# Transcriptomic-guided compound prioritization and proteomics validation for *HNRNPU* deficiency identify signalling correction

**DOI:** 10.64898/2026.05.04.722615

**Authors:** Xuan Ye, Diana Tikhomirova, Marika Oksanen, Massimiliano Gaetani, Hassan Gharibi, Francesca Mastropasqua, Kristiina Tammimies

## Abstract

*Heterogeneous nuclear ribonucleoprotein U (HNRNPU)* deficiency is a rare genetic cause of neurodevelopmental disorders (NDDs) lacking targeted therapies. Here, we developed a transcriptomic-guided compound prioritization pipeline using Connectivity Map (CMap) analysis on multi-model transcriptomic signatures from *HNRNPU*-deficient human cells and mouse models. Ten compounds were selected through manual curation and functionally screened in patient-derived *HNRNPU*-deficient neuroepithelial stem (NES) cells with earlier observed cellular phenotypes. Two of the compounds, AS601245 and Lenalidomide, significantly reduced the elevated neural progenitor population during differentiation, and their combination further decreased primary cilia incidence, indicating partial rescue of the patient-specific cellular phenotypes. To understand the mechanisms underlying the partial rescue, we employed proteome integral solubility alteration (PISA) and expression proteomics. PISA assay identified TMEM150C and GSK3A as proximal targets of combined treatment. Additionally, we observed reversal of multiple biological pathways including downregulation of Wnt signalling and upregulation of mitochondrial pathways and transmembrane proteins. Altogether, we established a computational-experimental pipeline for transcriptomic-guided drug repurposing for a monogenic NDD, and demonstrated that the network-level modulation partially rescues the delayed neural differentiation in *HNRNPU*-deficient neural cells.

## Introduction

Heterogeneous nuclear ribonucleoprotein U (HNRNPU), also known as Scaffold Attachment Factor A (SAF-A) [1], is an RNA-binding protein (RBP) ubiquitously expressed across multiple tissues, with the highest levels in the brain [2]. The HNRNPU protein is primarily located in the cell nucleus and acts as a nuclear chromatin organizer, mainly involved in transcription [3], alternative splicing [4–6], chromatin stabilization and remodelling [7–9]. More importantly, the *HNRNPU* gene has been linked to various neurodevelopmental disorders (NDDs). Pathogenic *de novo* heterozygous loss-of-function variants and deletions in *HNRNPU* locus cause an NDD (*HNRNPU*-related NDD), characterized by global developmental delay, intellectual disability, early-onset seizures and craniofacial features, and with variability for having autism and hypotonia [10–12].

Several studies, including our previous research, have demonstrated the essential role of HNRNPU in transcriptomic regulation during brain development. Mouse models of *Hnrnpu* haploinsufficiency showed transcriptomic changes in the hippocampus and neocortical cells [13]. Tissue-specific *Hnrnpu* loss of function in mouse models revealed that *HNRNPU* truncation caused variable alternative splicing alterations, leading to defects in mouse heart [6] and cortex [14, 15]. Complementing these findings, *in vitro* human models such as those using *HNRNPU*^+/−^ human cortical organoids exhibited transcriptome-wide dysregulation that overlapped with patterns observed in embryonic mouse models, showing disruptions in the neurodevelopmental process [16]. In our previous study, we developed an *in vitro* human *HNRNPU* locus deficiency model using patient-specific induced pluripotent stem cells (iPSCs)-derived neuroepithelial stem (NES) cells with a hindbrain profile [17]. We confirmed that *HNRNPU* deficiency caused chromatin remodelling, epigenetic changes and transcriptional rewiring during neural differentiation [17, 18]. At the cellular level, *HNRNPU* downregulation delayed commitment to neural stem cell differentiation, characterized by a higher proportion of neural progenitors during neuron maturation [17].

There is an unmet need for targeted pharmacological treatments for *HNRNPU*-related NDDs, despite recent research that has advanced in describing the mechanistic insights of *HNRNPU* in the brain. Because *HNRNPU* deficiency affects multiple regulatory pathways simultaneously, approaches that modulate global transcriptional patterns may offer an effective strategy to counteract these broad molecular consequences. Transcriptomic reversal methods use expression signatures to identify compounds that shift perturbed profiles toward a more typical state [13, 19]. The Connectivity Map (CMap)(clue.io) [20] database provides a large publicly available catalogue of compound-induced transcriptional responses, generated predominantly from cancer cell lines [21, 22]. Several studies have successfully used CMap-based signature matching to advance drug repurposing for cancer [23–25] and other non-neurological diseases [26–28]. Although some studies conceptualized the use of CMap in neurodevelopmental settings [13, 29], the following functional validation and mechanistic insights of the computational screening results remain largely unexplored.

In this study, we integrated differential expression signatures from human and mouse *HNRNPU* deficiency models to prioritize compounds predicted to counteract the transcriptional changes associated with its deficiency. We first tested phenotypic outcomes of these candidates in patient-derived NES cells. We then used two deep, quantitative proteome profiling approaches: proteome integral solubility alteration (PISA) assay [30] to identify proximal targets of the prioritized compounds and expression proteomics to identify molecular pathways affected by the compounds. Our combined computational and experimental framework aims to identify compounds that modulate *HNRNPU*-related developmental phenotypes and elucidate the early mechanisms associated with the transcriptional modulation affecting neural differentiation.

## Results

### Prioritization of test compounds for reversing *HNRNPU* deficiency transcriptomic signatures

We prioritized candidate compounds based on differentially expressed genes (DEGs) signatures from multiple published transcriptomic datasets associated with *HNRNPU* deficiency. As the primary dataset, we used bulk RNA sequencing data from HNRNPUdel NES cells compared with CTRL NES cells after 28 days of neural differentiation [17]. For cross-validation, we incorporated DEG signatures from additional datasets, including mouse hippocampal and neocortical datasets [13, 14] and human cortical organoids carrying *HNRNPU* variants [16]. For each dataset, we submitted pre-ranked up- and down-regulated genes to the CMap query platform [20] to identify compounds whose transcriptional profiles strongly opposed the *HNRNPU*-associated signatures. Compounds with connectivity scores ≤ −90 were considered strong candidates for reversing the expression pattern. From the list generated from DEGs for HNRNPUdel NES cells after 28 days of differentiation, we retained compounds that appeared in at least two of the four other datasets. Additionally, we excluded compounds known to inhibit cell proliferation or commercially unavailable, yielding ten suitable candidate compounds for downstream testing (**Table 1**). Among them, AG-1295, Lenalidomide, Ascorbyl-palmitate, and Lopinavir are negatively correlated to up-regulated genes, PSB-069, Physostigmine, PF-543, BRD6929, AS601245, and Mocetinostat are negatively correlated to down-regulated genes.

**Table 1.**
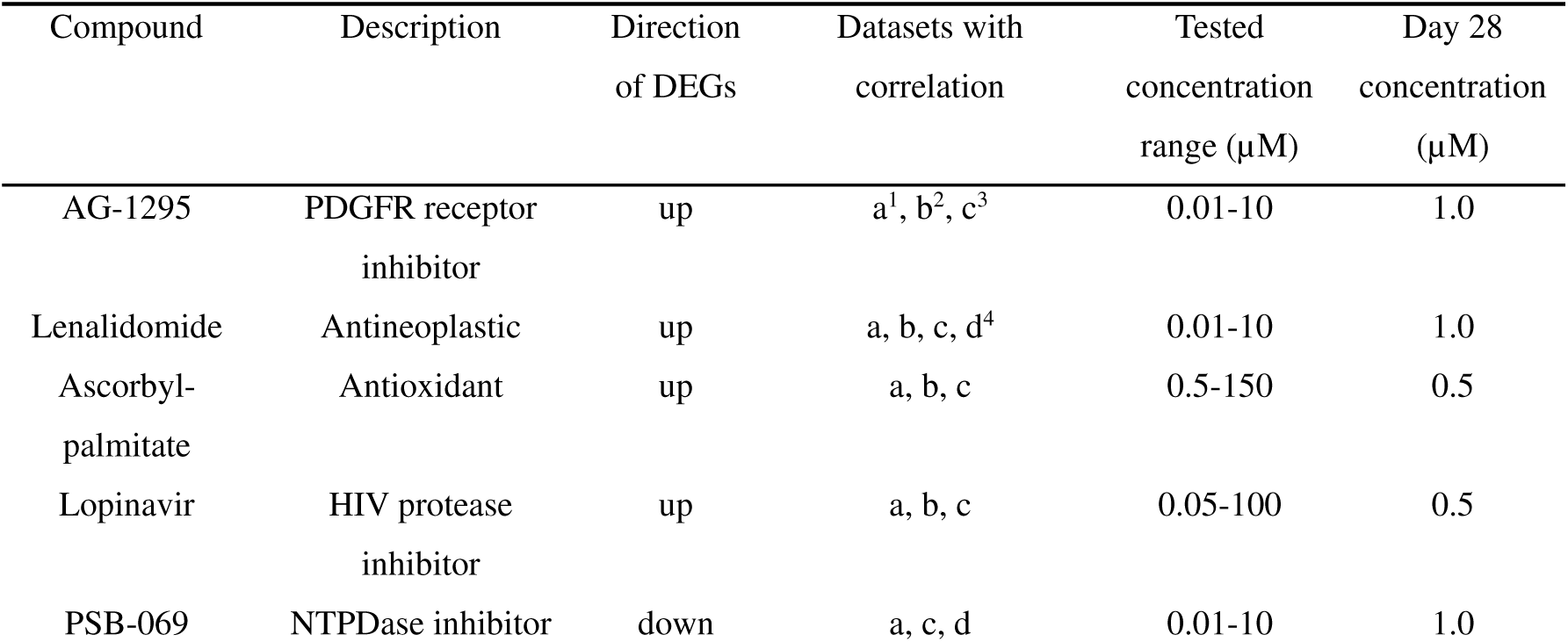
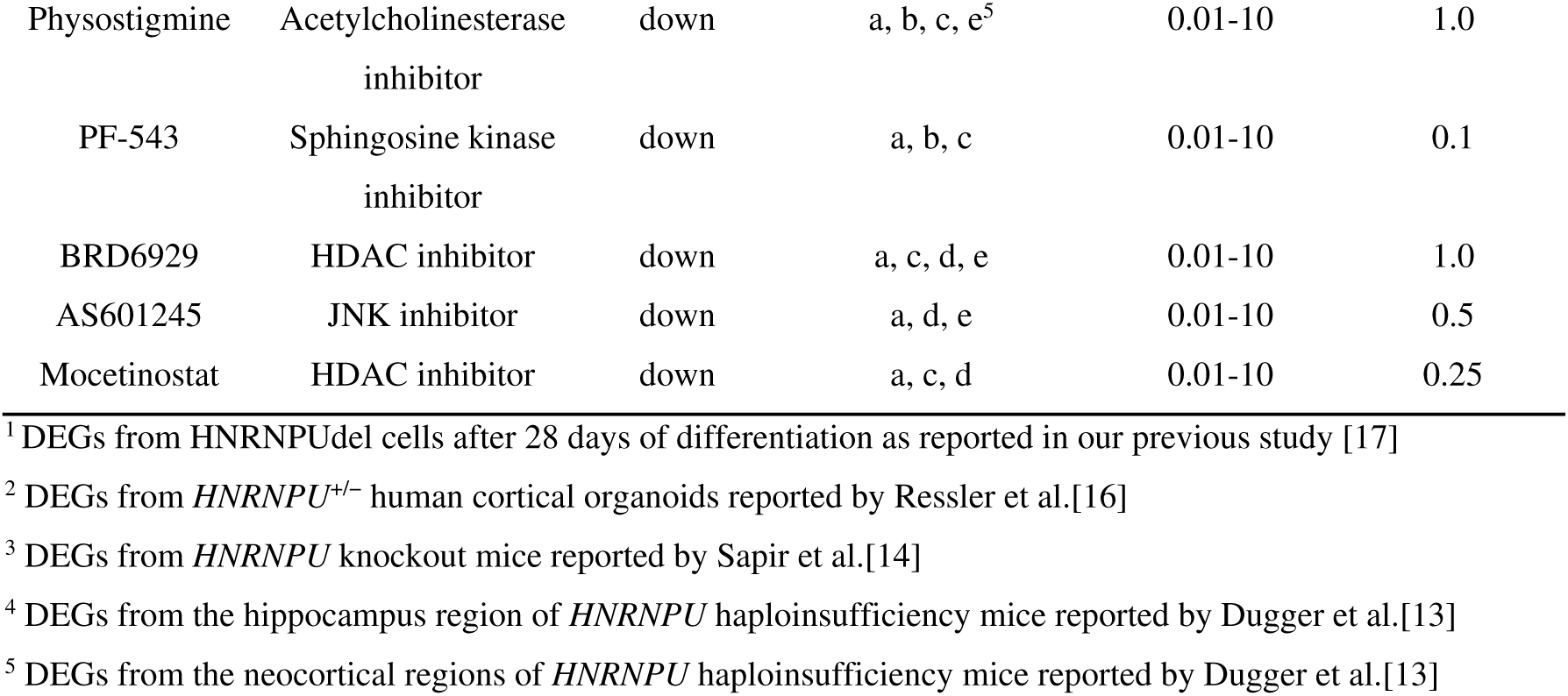
Selected test compounds based on CMap queries for the treatment of *HNRNPU* deficiency.

### AS601245 and Lenalidomide partially mitigate the excessive progenitor cell population in HNRNPUdel cells

To functionally evaluate the prioritized compounds, we utilized previously established iPSC-derived NES cell models with a hindbrain profile [17, 31, 32]. We included CTRL cells from a neurotypical male donor and HNRNPUdel cells from a male donor carrying a 44 kilobase (kb) heterozygous deletion spanning from the *COX20* to *HNRNPU* genes [33]. For 28-day (D28) studies, the NES cells were further differentiated for 28 days using an undirected protocol as previously described [34, 35]. During the differentiation, the multipotent neural progenitors progressively mature into neurons or other neural cell types, generating a mixed cell population of excitatory, inhibitory, and progenitor neural cells [35]. Our previous study delineated some key phenotypic changes in HNRNPUdel cells at D28 stage of neuronal differentiation, including increased proliferation rate, increased percentage of SOX2 (Sex determining region Y-box 2) positive cells, and increased percentage of ciliated cells [17], suggesting a delayed commitment to differentiation in HNRNPUdel cells.

Prior to long-term differentiation experiments, we performed dose-response assays to determine non-toxic concentration ranges for the ten selected compounds. NES cells were exposed to 16 concentrations of each compound for 24 and 48 hours, and viability was quantified by MTS [3-(4,5-dimethylthiazol-2-yl)-5-(3-carboxymethoxyphenyl)-2-(4-sulfophenyl)-2H-tetrazolium] assay (**Appendix Fig. S1**). Based on these data, we selected concentrations that maintained viability while maximizing potential efficacy (**Table 1**).

We next assessed whether the selected drug concentrations affected proliferation, as altered proliferation could confound the interpretation of differentiation outcomes. Using 5-bromo-2′-deoxyuridine (BrdU) incorporation assay, we confirmed that HNRNPUdel cells exhibited an elevated baseline proliferation rate compared to CTRL cells, both on day0 (D0) (P = 0.044, Student’s t-test) and D28 of differentiation (P = 0.016, Student’s t-test), consistent with our earlier findings [17] (**Appendix Fig. S2A-B**). Importantly, none of the compounds significantly changed the proliferation rate relative to non-treated (NT) HNRNPUdel cells, although AS601245, Lenalidomide and Lopinavir showed small downward trends (**Appendix Fig. S2B**). Due to pronounced cell death observed with Mocetinostat by day 8 of differentiation, data for Mocetinostat at D28 was not included. These results indicate that subsequent differentiation-related phenotypes cannot be explained by altered proliferation.

To assess whether the selected compounds could influence the earlier observed differentiation phenotype of HNRNPUdel cells, we quantified the proportion of SOX2-positive cells after 28 days of differentiation. We confirmed that HNRNPUdel cells had a larger proportion of SOX2 positive neural progenitor cells compared with CTRL at D28 stage (P = 5.15e-05, Student’s t-test) (**Fig. 1A**). Among the ten prioritized compounds, AS601245 and Lenalidomide treatment significantly reduced the percentage of SOX2-positive progenitors in HNRNPUdel cells (AS601245: P = 4.67e-05; Lenalidomide: P = 0.026, Student’s t-test) (**Fig. 1A**). These findings supported AS601245 and Lenalidomide as promising candidates for further evaluations. To examine whether the two compounds exert counteracting or additive effects, we included combination treatments: a high-dose combination (**Comb1**; 0.5 µM AS601245 and 1 µM Lenalidomide) and a low-dose combination (**Comb0.5**; 0.25 µM AS601245 and 0.5 µM Lenalidomide). As in the initial screen, none of the combination or single compound treatments significantly altered proliferation in *HNRNPU*del cells (**Fig. 1B**). However, all treatments significantly lowered the elevated proportion of SOX2-positive cells characteristic of HNRNPUdel cultures (**Fig. 1C-D**). AS601245 and the two combination treatment groups had more pronounced effects compared to Lenalidomide (**Fig. 1D**).

**Figure 1.**
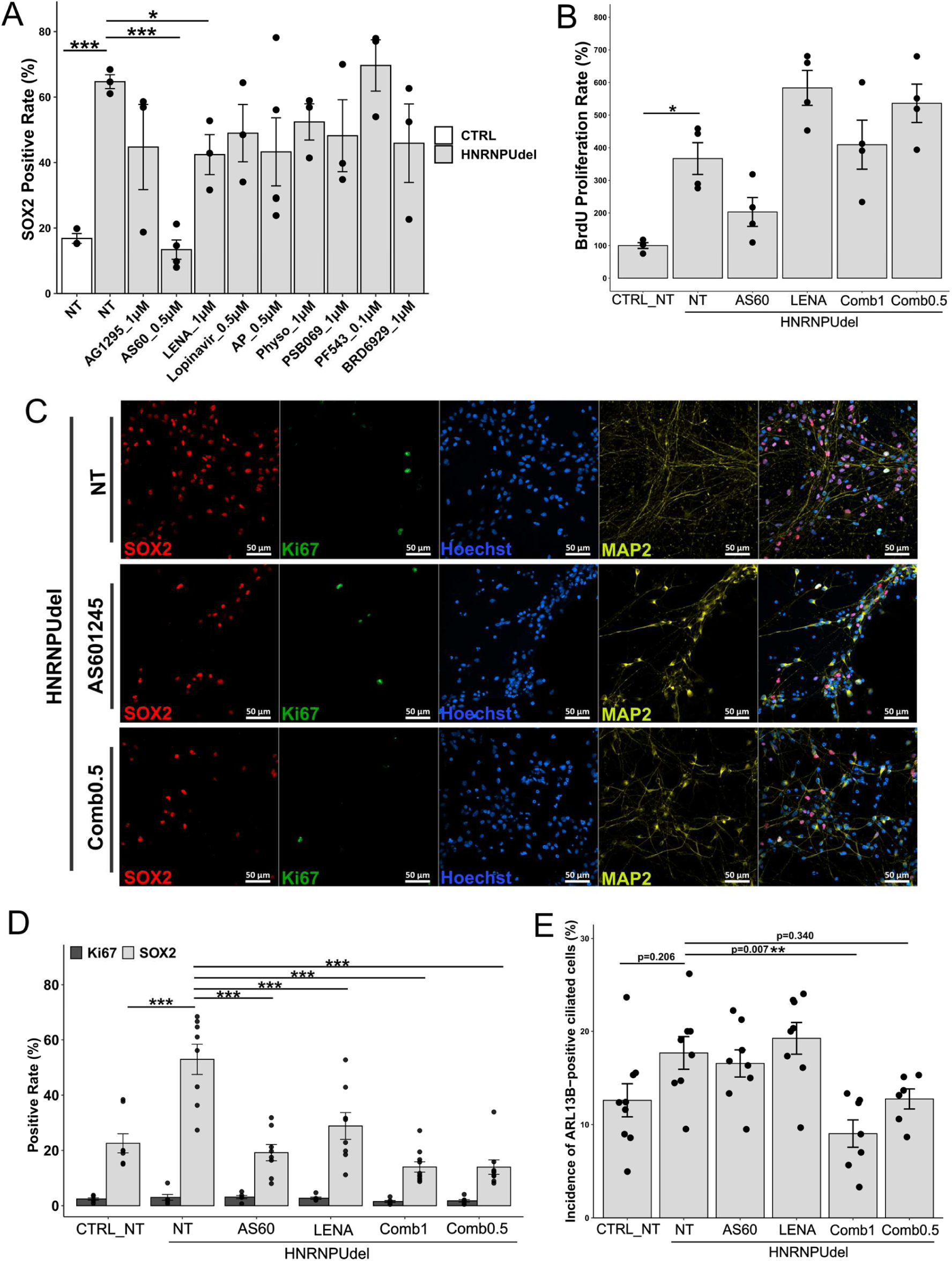
*HNRNPU* deficiency impairs neural differentiation and the phenotypic changes are partially rescued by AS601245 (AS60) and Lenalidomide (LENA). (A) Quantification of the percentage of SOX2-positive neural progenitor cells in CTRL and HNRNPUdel cultures under non-treated (NT) conditions and following treatment with nine prioritized compounds in HNRNPUdel cells. (B) Cell proliferation rates in CTRL and HNRNPUdel cells under NT conditions and after treatment with AS601245, Lenalidomide, **Comb1** (0.5 µM AS601245 + 1 µM Lenalidomide), and **Comb0.5** (0.25 µM AS601245 + 0.5 µM Lenalidomide). (C) Representative immunofluorescence images showing SOX2 (red), Ki67 (green) and MAP2 (yellow) staining in HNRNPUdel cells under NT, AS601245, and **Comb0.5** treatment; nuclei are counterstained with Hoechst (blue). Scale bar: 50 µm. (D) Quantification of SOX2- and Ki67-positive cells based on immunofluorescence analysis. (E) Quantification of ARL13B-positive primary ciliated cells based on immunofluorescence staining. Data are presented as mean ± SEM (n = 3–5 biological replicates). *p < 0.05, **p < 0.01, *** p < 0.001 (one-way ANOVA with Tukey’s HSD test or Student’s t-test, as indicated in Methods).

Persistent primary cilia have been linked to impaired exit from the progenitor state in neurodevelopmental contexts, and HNRNPUdel cells previously showed altered ciliogenesis [17, 36, 37]. We, therefore, analysed the percentage of ADP-ribosylation factor-like protein 13B (ARL13B)-positive primary ciliated cells after D28 differentiation. In HNRNPUdel cells, the proportion of primary ciliated cells showed a non-significant increase compared to CTRL cells (P = 0.206, one-way analysis of variance (ANOVA) and Tukey’s honestly significant difference (HSD) test). Neither single-compound treatment, nor the low-dose combination (**Comb0.5**), led to a reduction in primary cilia incidence (**Fig. 1E**). However, the high-dose combination (**Comb1**) significantly decreased the percentage of ciliated cells (**Fig. 1E**) (P = 0.007, ANOVA and Tukey’s HSD test), demonstrating a synergistic effect in ameliorating the ciliogenesis phenotype associated with delayed neural differentiation in HNRNPUdel cells.

### PISA identifies direct or proximal drug targets

Previous functional assays showed that AS601245 and Lenalidomide, particularly in combination, could partially rescue the delayed neural differentiation phenotype in HNRNPUdel cells by reducing the number of SOX2-positive neural progenitors and ARL13B-positive primary cilia. To investigate the molecular changes underlying these effects, we next examined compound-induced proteomic changes.

First, to identify direct drug targets, we employed the PISA assay, which detects drug-induced alterations in protein solubility as a proxy for ligand interaction or stability shifts [30, 38]. For this, HNRNPUdel cells were treated for 30 minutes in three treatment groups: non-treated, 1.5 µM AS601245, and 1.5 µM AS601245 + 3 µM Lenalidomide. Significant PISA signals (P < 0.05) revealed a set of proteins consistently modulated by both AS601245 alone and the combination treatment (**Fig. 2A, Appendix Fig. S4C–D**). TMEM150C, CA11, and GSK3A were upregulated and GRM8 was downregulated in both treatments (**Fig. 2A**). PRSS12 and TENT4A were specifically upregulated and ZNF780A downregulated in the AS601245 treatment. Conversely, IGF2 and HES1 were downregulated only in the combination group (**Fig. 2A**). Overall, the PISA results underscore TMEM150C, a regulator of mechanosensitive ion channels [39], as a direct target of AS601245, and suggest that drug exposure alters as first targets proteins involved in cell growth and differentiation (GSK3A, HES1, IGF2, EVC), as well as transcriptional regulation (BRF2, CA11, ZNF780A, KMT5B). Together, these changes point towards these drugs affecting convergent signalling pathways relevant for neurogenesis.

**Figure 2.**
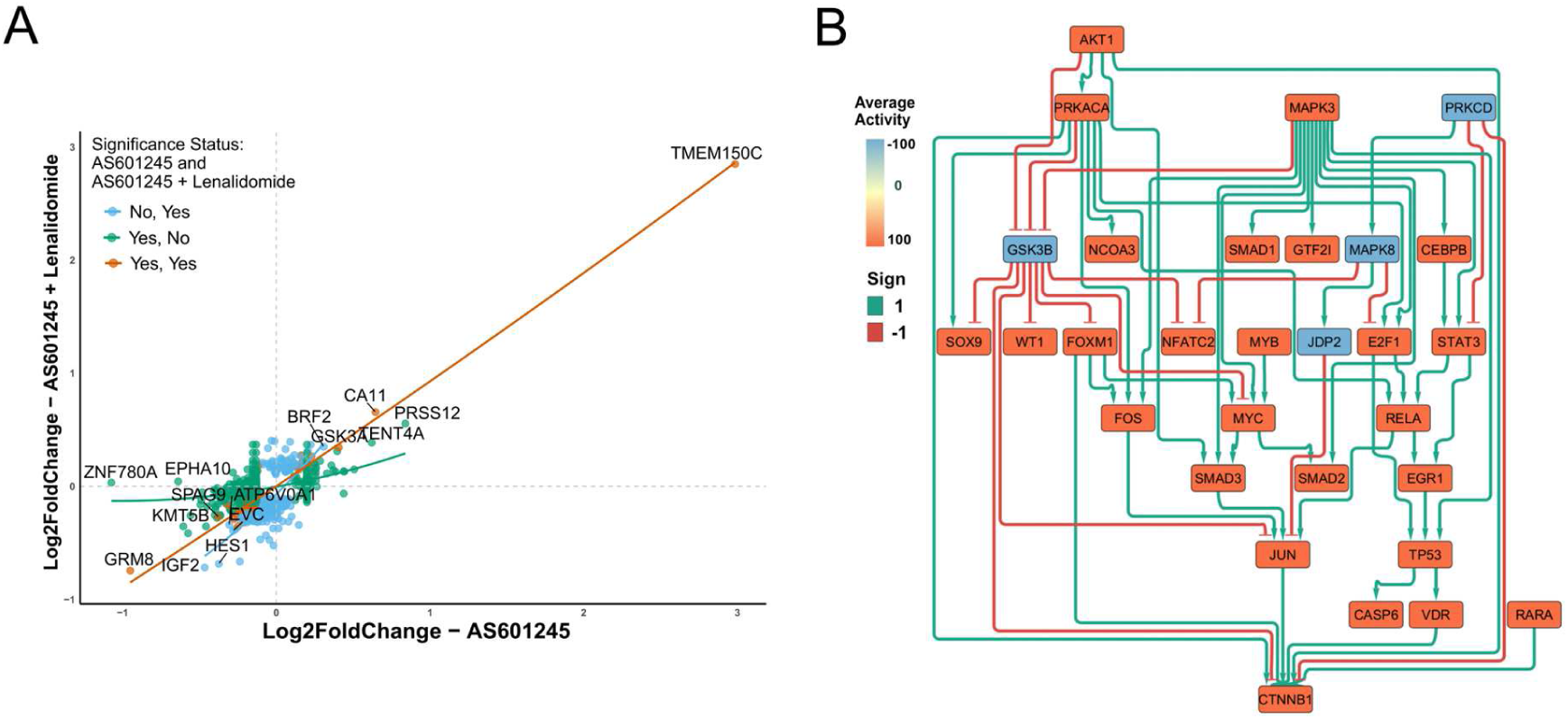
PISA and computational inference identify proximal drug targets and upstream networks. (A) Correlation plot from PISA assay showing Log2(Fold Change) values between single AS601245 treatment and combined treatment in HNRNPUdel cells. Significant proteins (unadjusted p < 0.05) are shown and highlighted by colour (orange: significant in both treatments; dark green: significant only in AS601245 treatment; sky blue: significant only in combined treatment). (B) Inferred causal networks from D5 transcriptomic signatures in HNRNPUdel cells. All nodes are included, with a betweenness centrality > 0.

In parallel, we integrated the earlier published longitudinal D0, day5 (D5) and D28 transcriptomic signatures from HNRNPUdel cells using causal network modelling to infer upstream regulatory interactions. The inferred networks highlighted kinase-associated signalling nodes, including MAPK1/3 and AKT1/GSK3B, together with PLK-related cell cycle pathways as prominent regulatory hubs at later differentiation stages (**Fig. 2B, Appendix Fig. S3A-B**). Notably, the emergence of GSK3 signalling in the inferred network is consistent with the detection of GSK3A among the proteins with altered solubility in the PISA assay, supporting the relevance of kinase-mediated signalling pathways in the observed phenotype and rescue.

### Proteomics reveals mechanisms of action underlying phenotypic rescue

To investigate global proteomic changes at D28 of differentiation and treatment, we extracted proteins from CTRL, non-treated (NT) HNRNPUdel, and **Comb1**-treated HNRNPUdel cells. In total, we quantified 10,279 proteins with at least two unique peptides after filtering out the contaminants. Principal component analysis (PCA) showed clear separation among the three groups (**Appendix Fig. S4A**), while normalized protein abundance distributions were consistent across samples, indicating robust data quality (**Appendix Fig. S4B**).

Given that *HNRNPU* deficiency disrupts genome-wide transcriptional regulation [17], and the candidate compounds were prioritized due to their potential to counteract these transcriptomic signatures, we first compared the global protein expression profiles in HNRNPUdel with our previously obtained RNA-sequencing data at the same time point (**Fig. 3A**). Most significantly altered proteins showed concordant directional changes (**Fig. 3A**), with a strong positive correlation between proteomics and RNA-sequencing datasets (Pearson r = 0.72, p < 2.2e-16), confirming that major transcriptomic perturbations extend to the proteomic level.

**Figure 3.**
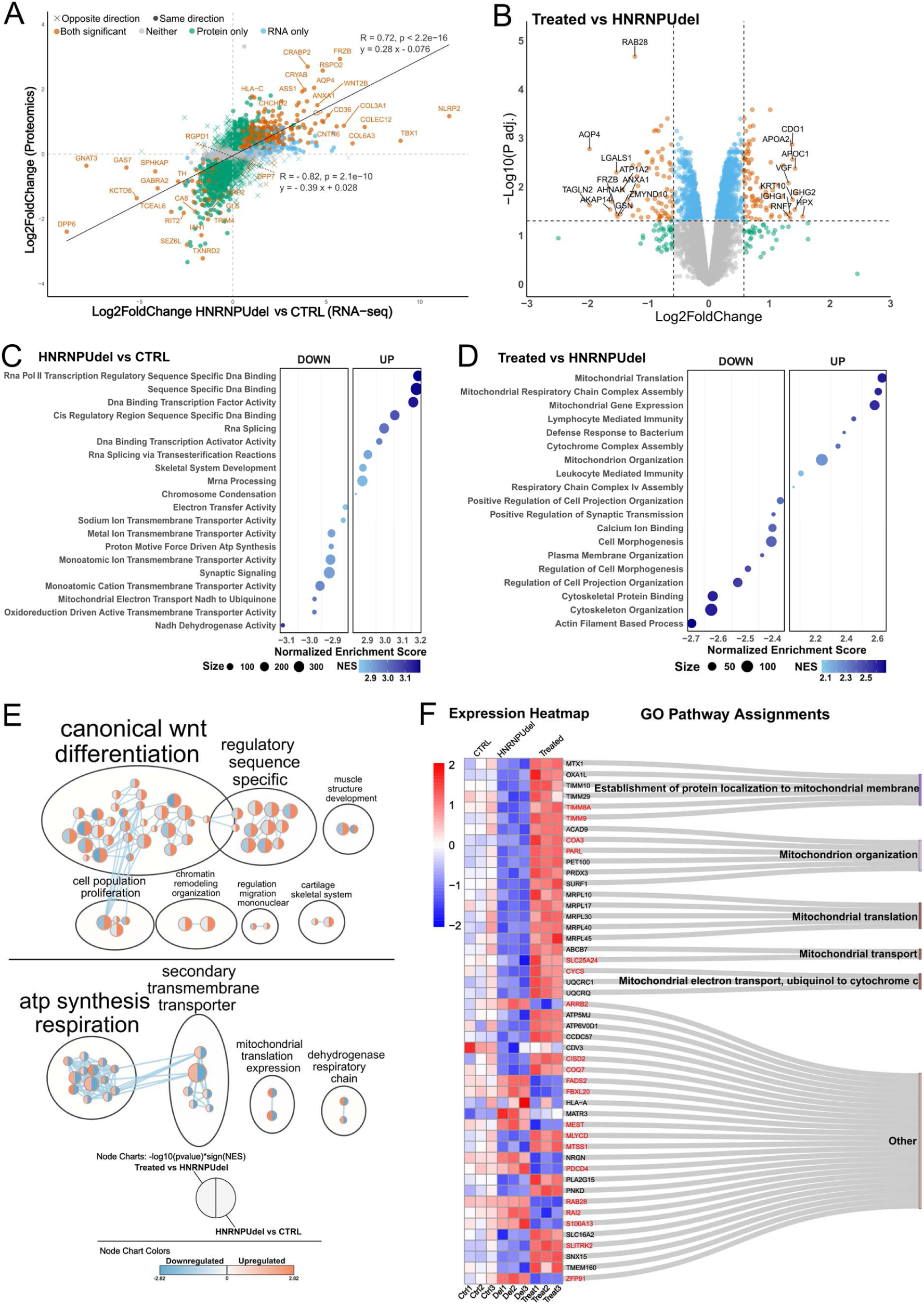
Proteomics profiling elucidates key modulators and pathways underlining partial rescue in HNRNPUdel cells after treatment. (A) Scatter plot of matched Log2(Fold Change) values comparing RNA-sequencing and proteomics data for D28 HNRNPUdel cells versus CTRL. Points represent individual genes/proteins and colours represent the significance (orange: significant in both datasets; dark green: significant only in proteomics; sky blue: significant only in RNA-sequencing; grey: significant in neither dataset). Correlation curves of the same directional changed points (dots and solid line) and opposite directional changes (crosses and dashed line) and correlation coefficients are shown. (B) Volcano plot depicting protein expression changes in HNRNPUdel cells after D28 combined drug treatment compared to non-treated group. Points are coloured by significance (significance thresholds: adjusted p value < 0.05, |Log2(Fold Change)| > 0.58). (C) Bubble plot of the top 10 upregulated and downregulated GO terms based on ranked differentially expressed proteins in HNRNPUdel cells versus CTRL after 28 days of differentiation. Bubble size denotes gene count; colour indicates normalized enrichment score. (D) Bubble plot of the top 10 upregulated and downregulated GO terms in HNRNPUdel cells following D28 combined treatment versus non-treated. Bubble size denotes gene count; colour indicates normalized enrichment score. (E) Selected enriched pathways showing the reversal effect at D28. For each node, the left half indicates the enrichment in HNRNPUdel combined treatment versus HNRNPUdel non-treated, and the right half indicates the enrichment in HNRNPUdel non-treated versus CTRL (orange: upregulated; blue: downregulated). (F) Heatmap of normalized protein expression levels demonstrating reversal effects at D28 across selected proteins in CTRL, HNRNPUdel NT, and HNRNPUdel **Comb1**-treated groups. Associated GO terms for protein subsets are annotated on the right. Protein labels in red font indicate concordant changes in the corresponding RNA-sequencing data in HNRNPUdel versus CTRL.

We next examined the global protein expression patterns in the samples. We observed 846 significantly more abundant proteins (adjusted p value < 0.05 Benjamini-Hochberg adjustment, |Log2FoldChange| > 0.58) between HNRNPUdel and CTRL cells (**Appendix Fig. S4F**), as well as 157 significantly more abundant proteins between HNRNPUdel treated and non-treated cells (**Fig. 3B**). Gene Set Enrichment Analysis (GSEA) of the pre-ranked differentially abundant proteins showed that pathways related to RNA binding and chromatin regulation were upregulated in HNRNPUdel cells in comparison with CTRL cells, whereas ion channels and mitochondria-related electron transport chain pathways were downregulated (**Fig. 3C**). Following the combined treatment, mitochondria-related transcription, translation, and respiratory chain were upregulated and pathways related to cytoskeleton, projection, and cell morphology were downregulated (**Fig. 3D**). To identify pathways most strongly associated with cellular phenotypic reversal, we filtered for categories showing opposite enrichment in non-treated HNRNPUdel versus CTRL and **Comb1**-treated versus non-treated HNRNPUdel cells. Pathways upregulated in HNRNPUdel cells but in turn downregulated by the treatment included Wnt signalling, cell differentiation, DNA/RNA binding and chromatin remodelling, and cell morphogenesis. Conversely, pathways downregulated in HNRNPUdel cells but upregulated by the treatment involved transmembrane transporters and mitochondrial-related pathways (**Fig. 3E**).

Additionally, we selected those proteins whose expression was altered in HNRNPUdel versus control and showed reversed differential abundance upon drug exposure. Gene Ontology (GO) enrichment analysis on this subset revealed a strong association with mitochondria structure and cellular energy metabolism (**Fig. 3F**). Components of ubiquitination pathways, including FBXL20 and ZFP91, showed increased expression in HNRNPUdel cells and were reduced after treatment (**Fig. 3F**), suggesting drug-induced restoration of protein-homeostasis mechanisms. Together, we show strong mRNA–protein concordance in HNRNPUdel cells and the effects of combined AS601245-Lenalidomide treatment on reversing multiple dysregulated pathways, including Wnt signalling and mitochondrial transcription, translation, and respiration. These findings suggest effects on mitochondrial pathways and, modulation of developmental signalling pathways as potential contributors to the observed cellular phenotypic reversal.

## Discussion

*HNRNPU*-related NDDs arise from haploinsufficiency of a chromatin-associated RNA-binding protein that regulates transcription, alternative splicing and genome architecture among other roles [4–6, 40–42]. Consequently, *HNRNPU* deficiency induces coordinated network-level perturbations during neuronal development rather than disrupting a single downstream pathway [18]. Although mechanistic understanding has expanded through mouse models [13, 14], cortical organoids [16], and patient-derived neural stem cell systems [17], targeted therapeutic interventions remain largely unexplored. Here, we applied a transcriptomic reversal framework using CMap [21, 22] to prioritize compounds predicted to counteract *HNRNPU*-deficiency-associated molecular signatures. By this approach, we identified JNK inhibitor (AS601245) [43, 44] and immunomodulatory agent Lenalidomide [45] as candidate compounds capable of partially correcting the delayed neural differentiation and omics signatures identified in patient-derived neural cells.

Functionally, AS601245 and Lenalidomide reduced the elevated proportion of SOX2-positive progenitors observed in HNRNPUdel cells without significantly altering proliferation rates. Furthermore, the reduction in primary cilia incidence following the combination treatment further supports partial restoration of the differentiation dynamics in HNRNPUdel cells. Given the central role of primary cilia in coordinating Wnt and Hedgehog signalling during neural fate decisions [17, 36, 37, 46, 47], these findings suggest that pharmacological treatment applied here rebalances developmental signalling networks altered in the *HNRNPU* deficiency state.

We also provided additional mechanistic insights into the rescue using proteomics. The strong concordance between transcriptomics and proteomic signatures confirmed that *HNRNPU* deficiency induces stable downstream protein-level perturbations, validating the model as a translational platform. The PISA proteomic profiling [30, 38], identified TMEM150C, and GSK3A as primary key proteins directly modulated by drug exposure. As a member of the Damage-Regulated Autophagy Modulator (DRAM) family, TMEM150C modulates autophagic flux, cell survival, and mechanosensitive ion channels, potentially influencing apoptosis via JNK-dependent pathways [39, 48–50]. Given the known JNK inhibition by AS601245 [43, 44], the regulation of TMEM150C may suppress pro-apoptotic signalling, promoting progenitor commitment to neurogenesis [51]. GSK3 is a component of the Akt signalling pathway and takes part in the mTOR signalling during synaptic stimulation [52]. GSK3 is central to multiple intracellular pathways, including those activated by Wnt/β-catenin, Notch, Sonic Hedgehog [53, 54], thereby regulating crucial cellular functions, including cell proliferation, differentiation, and neural signalling. Furthermore, re-analysis of our earlier RNA-sequencing data using causal network analysis revealed GSK-signalling as one of the upstream key regulators. JNK inhibition may disrupt stress-induced signalling, thereby indirectly enhancing GSK3 activity and influencing downstream targets such as c-Jun [55]. The stabilized GSK3 could reinforce β-catenin degradation, suppressing Wnt-mediated transcription [54], aligning with Wnt signalling modulation in global proteomics. Additionally, PISA identified IGF2 as possible target of the combined treatment. The interplay of GSK3/IGF2 has been shown to improve mouse cognitive behaviour [56], suggesting IGF2 as one of the downstream proteins following GSK3 normalization. Considering IGF2 regulates neural stem cell proliferation through the PI3K/Akt pathway [57], the PISA results suggest a boarder PI3K/Akt/mTOR signalling pathway modulation triggered by the AS601245-Lenalidomide treatment.

Additionally, in this study we highlighted altered mitochondrial pathways as additional cellular phenotypes of *HNRNPU* deficiency, as we identified downregulation of key proteins of the electron transport chain, oxidative phosphorylation complexes, and mitochondrial translation machinery. Those enriched pathways were substantially reversed by combined AS601245 and Lenalidomide treatment, suggesting a possible restoration of bioenergetic pathways essential for neural differentiation [58]. Additionally, ion transmembrane transporters and other ion channel pathways were downregulated following treatment, indicating a rescue of the ion channel dysfunction involved in neurodevelopmental disorders, especially in epilepsies [59–61]. The coordinated modulation of ion transport and membrane-associated pathways further indicates integrated regulation of cellular energetics and signalling rather than isolated pathway correction.

Lenalidomide, an immunomodulatory agent targeting cereblon-dependent ubiquitination pathways, is best characterized in hematologic contexts but also modulates cytokine signalling and transcriptional regulators [62, 63]. Immune and inflammatory signalling have been increasingly implicated in neurodevelopmental trajectories, and exploratory studies have reported behavioural improvement in subsets of autistic individuals with elevated inflammatory markers following Lenalidomide exposure [64]. In our system, Lenalidomide alone produced moderate effects but enhanced AS601245-mediated rescue, particularly in relation to ciliogenesis, suggesting convergence between kinase-dependent signalling and ubiquitination-mediated regulatory pathways. The partial nature of the rescue is consistent with the expectation that chromatin-associated *HNRNPU* haploinsufficiency produces widespread regulatory imbalance [17], unlikely to be fully normalized by single-agent intervention.

The translational considerations for the two identified compounds require careful evaluation. AS601245 is not clinically approved, and its pharmacokinetics and brain penetrance in developmental contexts remain to be established. Lenalidomide is clinically approved but carries known teratogenic and immunomodulatory risks [65], necessitating stringent safety evaluation in paediatric settings. Furthermore, *in vitro* concentrations may not directly translate to achievable central nervous system exposure. Future work should therefore prioritize validation in additional genetically independent *HNRNPU* models, single-cell transcriptomic assessment of post-treatment state transitions, and *in vivo* pharmacodynamic studies in *Hnrnpu* haploinsufficient animals.

Several other limitations of this study should be acknowledged. First, the CMap database is heavily skewed toward cancer cell lines, leading to potential mismatches in pathway dynamics between oncogenic processes and neurodevelopmental contexts. Therefore, the functional validation in neurological models is required for screening drug candidates. Second, changes in protein abundances and solubility can describe two different proteomics dimensions, but dynamic signalling events might require additional phosphoproteomics to describe kinase-activated pathways, which might be relevant to decipher drug-induced modifications. Additionally, our reliance on a single patient-derived cell line and a neurotypical control restricts generalizability. Hence, the validation in multiple lines with heterogeneous *HNRNPU* mutations is crucial for robustness.

In summary, we established a computational-experimental pipeline for transcriptomic-guided drug repurposing in monogenic NDD characterized by disrupted regulatory networks. By integrating multi-model gene expression data, CMap-based prioritization, quantitative cellular phenotyping, and proteomic validation, we demonstrate that network-level pharmacological modulation can partially restore developmental progression in *HNRNPU*-deficient neuronal cells. This framework provides a scalable precision medicine strategy for additional chromatin-and RNA-binding protein–associated NDDs in which transcriptional dysregulation represents a central pathogenic mechanism.

## Methods

### Cell culturing and differentiation

The NES cells used in this study were generated by dual-SMAD inhibition of human induced pluripotent stem cells (iPSCs) derived from donors’ skin fibroblasts as described before [17, 31, 32]. The cell lines include cells from a neurotypical male donor (hereafter, CTRL) [31, 32], and cells from a male donor carrying a 44 kilobase (kb) heterozygous deletion spanning from *COX20* to *HNRNPU* genes (hereafter, HNRNPUdel), characterized earlier [33]. Informed consent from the donors were obtained earlier accordance to Swedish Ethical approval. NES cells were cultured as previously described [17]. In short, the NES cells were seeded on plastic culture ware, precoated with 20 µg/ml poly-L-ornithine (Sigma Aldrich), and 1 µg/ml laminin (Sigma Aldrich), and cultured in DMEM/F12 + Glutamax medium (Gibco) supplemented with 0.05× B-27 (Gibco), 1× N-2 (Gibco), 10 ng/ml bFGF (Life Technologies), 10 ng/ml EGF (PeproTech) and 10 U/ml penicillin/streptomycin (Gibco), maintained in 5% CO_2_ atmosphere at 37°C. The NES growth medium was changed daily. For immunofluorescence, the glass coverslips were precoated with 100 µg/ml poly-L-ornithine and 2 µg/ml laminin. For neural differentiation, the NES growth medium was changed to differentiation medium, containing 0.5× B-27,1× N-2 and 0.4 µg/ml laminin, without bFGF and EGF. Half of the differentiation media was changed every second day.

### Compounds prioritization by Connectivity Map

We used DEGs derived from multiple transcriptomic datasets related to *HNRNPU* deficiency to prioritize candidate compounds. The primary dataset is bulk RNA sequencing data from HNRNPUdel NES cells compared to CTRL NES cells after 28 days of differentiation [17]. For cross-validation, we incorporated additional DEGs from independent sources: a) single-cell RNA sequencing data of the hippocampal and neocortical regions, respectively, from *Hnrnpu* haploinsufficient mice as described by Dugger et al. [13]; b) RNA sequencing data of cortices derived from E13 conditional *Hnrnpu* knockout mice embryos, in Sapir et al [14]; c) single-cell RNA sequencing data of 45-day-old HNRNPU^+/−^ human cortical organoids in Ressler et al. [16].

Then, CMap (clue.io) [20] was used to screen and select the candidate compounds. From the transcriptomic datasets, we selected the significant DEGs (adjusted p value < 0.05) and ranked them by descending Log2 (Fold Change). We then added the top 150 up- and down-regulated DEGs into CMap as separate queries. The L1000 dataset (version 1.0) from the CMap was selected as the reference data for gene expression. Connectivity scores were derived from the enrichment of submitted genes in the reference gene sets caused by certain perturbations, ranging from -100 to 100. A positive score indicates a positive correlation between submitted genes and perturbations, and a negative score indicates a negative correlation. We selected compounds with connectivity scores less than -90, which are most likely to reverse the input gene signatures. We respectively conducted the CMap queries for above mentioned independent datasets and systematically compared the generated compound lists. From the compound list derived from HNRNPUdel NES cells after 28 days of differentiation, we screened compounds that appeared at least twice in the lists derived from the other four datasets. Subsequently, we excluded drugs known to inhibit cell proliferation and those that were commercially unavailable.

### Compounds information and preparation

AS601245 (CAS No. 345987-15-7), Lenalidomide (CAS No. 191732-72-6), ascorbyl palmitate (CAS No. 137-66-6), lopinavir (CAS No. 192725-17-0), AG-1295 (CAS No. 71897-07-9), physostigmine (CAS No. 57-64-7), PF-543 (CAS No. 1706522-79-3), BRD6929 (CAS No. 849234-64-6), mocetinostat (CAS No. 726169-73-9) were supplied by Nordic Biosite. PSB 069 (CAS No. 78510-31-3) was purchased from Bio-Techne^®^ Tocris. All drugs were supplied as powder or DMSO solution. The drugs in powder form were dissolved in an appropriate volume of DMSO, and all drugs were prepared as DMSO working solutions at final concentrations of 1 mM and 10 mM. Prior to the cell viability assay, the drugs were dispensed for 16 concentration gradients in logarithmic distribution (from 0.01 µM to 100 µM) in 384-well plates using Tecan^®^ D300e digital dispenser, and culture medium was added to a total volume of 40 µl/well. In all other cases, the drugs were diluted in culture medium to an appropriate concentration and added to the cells along with the medium. For combined treatment, **Comb1** refers to a combination of 0.5 µM AS601245 and 1 µM Lenalidomide mixed in cell media. **Comb0.5** refers to a combination of 0.25 µM AS601245 and 0.5 µM Lenalidomide in cell media.

### Cell viability assay

The NES cells were seeded in precoated 384-well plates (5,000 cells/well) and maintained for 24 hours prior to the treatment. Then the growth medium was changed to a previously prepared drug medium mix to a final volume of 25 µl/well. The drug medium mix was changed after 24 hours. NES cells treated with 0.1% DMSO were used as non-treated (NT) controls and defined as 100% viability. Cell viability was tested after 24 and 48-hour treatments. The CellTiter 96^®^ AQueous One Solution (Promega) was used to determine the number of viable cells. The CellTiter solution was added directly to the culture medium and incubated at 37°C for 3 hours, at 5% CO_2_ atmosphere. The absorbance was recorded at 490 nm using Tecan Infinite^®^ 200 PRO microplate reader.

### Cell proliferation assay

The NES cells were seeded, in 96-well plates (12,000 cells/well) for three replicates per condition, the day before the drug treatment. After 24 hours, the culture medium was changed to a differentiating drug medium mix and maintained for 24 hours, as well as 28 days for the BrdU assay. The BrdU-based proliferation assay using an ELISA kit (ab126556, Abcam) was performed to evaluate cellular proliferation difference. For this, 1× BrdU reagent was added to the cell media and incubated for 24 hours, allowing BrdU to incorporate into proliferating cells. Then the cell media was aspirated, and cells were fixed with the supplied fixing solution. After one wash with 1× plate wash buffer, the anti-BrdU monoclonal detector antibody was added and incubated for 1 hour at room temperature. The cells were then incubated with the horseradish peroxidase (HRP)-tagged secondary antibody for 30 minutes at room temperature. After the final water wash, the cells were exposed to TMB (3,3’, 5,5’-tetramethylbenzidine) substrate in the dark and the absorbance was recorded at a dual wavelength of 450/550 nm using Tecan Infinite^®^ 200 PRO microplate reader. To calibrate the effect of the number of nuclei in each sample, the cells were further incubated in TBS, 0.1% Triton-X 100, and Hoechst 33342 (Invitrogen) 1:1000 for 15 min, and the fluorescence intensity was measured. The proliferation rate of CTRL NT cells was defined as 100% and the statistical analyses were performed in R by unpaired Student’s *t*-test, or one-way analysis of variance (ANOVA) followed by the Tukey’s honestly significant difference (HSD) test.

### Immunofluorescence and image analysis

The NES cells were seeded in precoated glass coverslips (72,800 cells/well) the day before treatment. Then the growth medium was changed to differentiation medium together with drug compounds. The cells were maintained in drug medium mix and differentiated for 28 days prior to immunofluorescence analysis. Thereafter, the cells on coverslips were fixed with 4% paraformaldehyde for 20 minutes at room temperature, followed by four times 1× TBS wash. The blocking buffer was prepared using 5% Donkey Serum and 0.1% Triton X-100 in 1× TBS and was incubated with cells for 1 hour. The following primary antibodies were used: anti-SOX2 (Sex determining region Y-box 2) (ab5603); anti-Ki67 (#14-5698-82); anti-MAP2 (M2320); anti- PCNT (Pericentrin) (ab28144); anti-ARL13B (ADP-ribosylation factor-like protein 13B) (17711-1-AP). The primary antibodies were incubated overnight at +4°C after being diluted in the blocking buffer (SOX2 1:1000, Ki67 1:500, MAP2 1:1000, ARL13B 1:10000, PCNT 1:250). After four washing with 1× TBS, secondary antibodies (Alexa Fluor^®^ 488, Alexa Fluor^®^ 555, Alexa Fluor^®^ 633) were diluted 1:2000 in blocking buffer and incubated at room temperature for 1 h, protected from light. After three washes, Hoechst 33342 (Invitrogen) was diluted 1:2000 in blocking buffer and incubated for 15 minutes in the dark. Coverslips were then mounted with Diamond Antifade Mountant (ThermoFisher Scientific). Images were acquired with LSM 700 and LSM 900 Zeiss Confocal Microscope using 20× or 63× magnification.

For SOX2-positive cells analysis, at least three technical replicates from two biological replicates were analysed. For Ki67-positive cells, at least five technical replicates were analysed. For primary cilia analysis, two biological replicates of **Comb1** and **Comb0.5**, and three biological replicates of all the other treatments were analysed. For each replicate, at least three different areas were acquired and processed. The images were further analysed in ImageJ software (ver. 1.54f). The SOX2-positive, Ki67-positive and total nuclei were counted using an in-house macro, then the SOX2- and Ki67-positive rate was calculated accordingly. For primary cilia analysis, ARL13B-positive cells were considered ciliated and CiliaQ plugin [66] was used to identify and quantify the number of primary cilia. The statistical analyses were performed in R by one-way ANOVA, followed by the Tukey’s HSD test.

### Expression proteomics and PISA assay

The proteome integral solubility alteration (PISA) assay and expression proteomics were carried out independently and analysed separately but combined into one tandem mass tag (TMT)-based 18-plex experiment split in two to host both experiments in one LC-MS/MS analysis, with a total of three conditions in triplicate for PISA and three triplicates for expression proteomics.

For PISA assay, HNRNPUdel cells were differentiated for 2 days and treated in complete medium for 30min with DMSO as untreated control, with 1.5µM AS601245, or with 1.5µM AS601245 and 3µM Lenalidomide. Samples were processed according to the proof-of-principle PISA method [30] and the latest optimized protocols for higher proteome depth of analysis, extended TMT multiplicity and improved sensitivity [38, 67]. Briefly, cells were detached, washed twice in PBS, resuspended into PBS with Halt protease inhibitors (Halt protease inhibitors, Thermo Fisher), up to the final ultracentrifugation step of the soluble fraction after application of the protein precipitation gradient, pooling of temperature points for each samples, protein extraction by five freeze-thaw cycles in liquid nitrogen and 37°C with 0.4% NP40, and isolation of soluble proteins by ultracentrifugation.

For expression proteomics, the NES cells were differentiated for 28 days as described above and washed three times with PBS buffer. Three conditions were analysed: CTRL9 non-treated (DMSO control); HNRNPUdel non-treated (DMSO control); HNRNPUdel treated with combined 0.5µM AS601245 and 1µM Lenalidomide. Cells were then collected, washed twice in PBS and resuspended in radioimmunoprecipitation assay (RIPA) buffer with protease and phosphatase inhibitors. Cells were lysed by subjecting them to freeze-thaw cycles in liquid nitrogen for a total of 5 times, followed by probe sonication for 1 min (03 pulse, 03 pause, 10 times -30% amplitude) on ice. After centrifugation for debris removal at 14,000 at 4 °C for 30 min, the supernatant was collected.

The total protein concentration of extracted samples from PISA and expression proteomics was measured using micro-BCA (bicinchoninic acid) assay and 50 μM of protein was processed by 8 mM dithiothreitol (Sigma) reduction, 25 mM iodoacetamide (Sigma) alkylation. Then samples were precipitated using cold acetone at -20 °C overnight. Briefly, as previously published in the above-mentioned references, 50 µg of each sample was reduced, alkylated and precipitated using cold acetone. Samples were resuspended in EPPS (4-(2-Hydroxyethyl)-1-piperazinepropanesulfonic acid) buffer, 8 M urea, pH 8.0, diluted down to 4 M urea and digested by LysC, then diluted down to 1 M urea and digested with trypsin. Each digest was labelled using TMTpro 18-plex technology (Thermo Fischer) and a final multiplex sample was first cleaned by Sep-Pack C18 column (Waters). The final sample was fractionated off-line by capillary reversed phase chromatography at pH 10 into 48 fractions, and each of them was then analysed by high-resolution nLC–ESI-MS/MS (nanoscale liquid chromatography-electrospray ionization-tandem mass spectrometry), as described below. Then LC-MS/MS data were subjected to peptide/protein identification and quantification, and data analysis.

NanoLC-MS/MS analyses were performed on an Orbitrap Exploris 480 mass spectrometer (Thermo Fisher Scientific). The instrument was equipped with an EASY ElectroSpray source and connected online to an Ultimate 3000 nanoflow UPLC system. The samples were pre-concentrated and desalted online using a PepMap C18 nano-trap column (length - 2 cm; inner diameter - 75 µm; particle size - 3 µm; pore size - 100 Å; Thermo Fisher Scientific) with a flow rate of 3 µL/min for 5 min. Peptide separation was performed on an EASY-Spray C18 reversed- phase nano-LC column (Acclaim PepMap RSLC; length - 50 cm; inner diameter - 2 µm; particle size - 2 µm; pore size – 100 Å; Thermo Scientific) at 55 °C and a flow rate of 300 nL/min. Peptides were separated using a binary solvent system consisting of 0.1% (v/v) FA, 2% (v/v) ACN (solvent A) and 98% ACN (v/v), 0.1% (v/v) FA (solvent B). They were eluted with a gradient of 3–26% B in 97 min, and 26–95% B in 9 min. Subsequently, the analytical column was washed with 95% B for 5 min before re-equilibration with 3% B. The mass spectrometer was operated in a data-dependent acquisition mode. A survey mass spectrum (from m/z 375 to 1500) was acquired in the Orbitrap analyser at a nominal resolution of 120,000. The automatic gain control (AGC) target was set as 100% standard, with the maximum injection time of 50 ms. The most abundant ions in charge states 2+ to 7+ were isolated in a 3 s cycle, fragmented using HCD MS/MS with 33% normalized collision energy, and detected in the Orbitrap analyser at a nominal mass resolution of 50,000. The AGC target for MS/MS was set as 250% standard with a maximum injection time of 100 ms, whereas dynamic exclusion was set to 45 s with a 10-ppm mass window.

Proteome Discoverer 3.1 was utilized for the database search and quantification against the Uniprot Homo sapiens (Human) protein database UP000005640. Cysteine carbamidomethylation was set as a fixed modification, along with TMT-related modifications, deamination on methionine, oxidation on arginine, and asparagine as variable modifications. Enzyme specificity was defined as trypsin with a maximum of two missed cleavages. A 1% false discovery rate was employed as a filter at both the protein and peptide levels. Contaminants and reversed-hit peptides were removed, and only proteins with at least two unique peptides were included in the quantitative analysis. Proteins with missing values in all replicates for any treatment were eliminated. The quantified abundance of each protein in each sample (labelled with a different TMT) was normalized to the total intensity of all proteins in that sample. For each protein in each compound replicate, the normalized protein abundance was divided by the average abundance of that protein in the vehicle-treated replicates. The average ratio across replicates of each compound compared to the vehicle control was calculated, and the Log2 values of these ratios were determined. Principal component analysis (PCA) and normalized protein abundance distributions were performed for quality control. Differential expression analysis of the normalized protein abundance was proceeded using the *limma* package in R (v4.4.2) [68]. Differentially expressed proteins were selected compared to vehicle treated controls. The significance thresholds for expression proteomics data were adjusted p value < 0.05 (Benjamini-Hochberg adjustment), and absolute Log2 Fold Change > 0.58. The significance thresholds for PISA results were unadjusted p-value < 0.05 and absolute Log2 Fold Change > 0.25.

### Pathway enrichment analysis

Pathway enrichment analysis was done in reference to the previously described protocols [69]. In short, a ranked list of significantly differentially abundant proteins was used in Gene Set Enrichment Analysis (GSEA) software (v 4.4.0) to analyse the enrichment in the gene set database Gene Ontology Biological Process (GO BP) and Gene Ontology Molecular Function (GO MF). The GSEA reports were visualized in R (v4.4.2) as bubble plots. Additionally, the GSEA output was visualized in Cytoscape (v3.10.4) with Enrichment Map and AutoAnnotate [69–71] after selecting the nodes that have opposite regulation directions between “HNRNPUdel vs CTRL” group and “HNRNPUdel treated vs HNRNPUdel non-treated” group.

For the expression heatmap, proteins with significant (adjusted p.value < 0.05, Benjamini-Hochberg adjustment) and opposite directions of changes were selected. GO analysis was performed on this subset of proteins by clusterProfiler [72] and assigned to the protein list. Unassigned proteins were grouped into “Other”. The heatmap was generated from the Log2 abundance values from the normalized data and visualized in R (v4.4.2).

### Causal signalling network inference from transcriptomic data

Causal networks were inferred using previously generated RNA-sequencing data from HNRNPUdel NES cells at differentiation days 0, 5 and 28 [17]. Pathway activities were estimated using PROGENy (v1.32.0) [73] and transcription factor activity using the decoupleR framework (v2.16.0) [74] from the gene expression footprints. To minimise the complexity of the network, the top 100 most active transcription factors serve as the downstream targets. Pathway activity scores were used to weigh the optimisation. The integer linear programming (ILP) problem was solved using the IBM CPLEX solver (v22.1.1) with a time limit of 900 seconds and a relative mixed-integer programming (MIP) gap tolerance of 0.05. These activity scores were integrated as inputs to the CARNIVAL pipeline [75] to reconstruct the most parsimonious causal signalling network based on a signed and directed prior knowledge network generated from the OmniPath (v3.19.1) database [76]. The resulting optimised networks were pruned to remove inactive edges (weight = 0) and isolated nodes. The networks were visualised in Cytoscape (v3.10.4) and used to identify high-centrality regulatory hubs.

## Data availability

The mass spectrometry proteomics data have been deposited to the ProteomeXchange Consortium via the PRIDE [77] partner repository with the dataset identifier PXD076404. Code used for the analyses in here are available in GitHub https://github.com/Tammimies-Lab/Transcriptomics_drug_re_HNRNPU

Further information and requests for resources and reagents should be directed to and fulfilled by the Lead Contact, Kristiina Tammimies.

## Author contributions

Conceptualization: X.Y., F.M., and K.T.; Methodology and Data Analysis: X.Y., D.T., M.O., M.G., H.G., F.M. and K.T.; Validation: X.Y., M.O. and K.T.; Resources: M.G. and K.T.; Data Curation: X.Y., H.G., and M.G.; Writing – Original Draft: X.Y. and K.T.; Writing – Review & Editing: X.Y., F.M., and K.T.; Visualization: X.Y., D.T.; Supervision: F.M. and K.T.; Project Administration: K.T.; Funding Acquisition: K.T., Critical review of the draft: All authors.

## Acknowledgement

We thank Chemical Proteomics at Karolinska Institutet (Chemistry I Division, MBB Department), unit of SciLifeLab and node of the Swedish National Infrastructure for Biological Mass Spectrometry (BioMS), for providing full support in the experimental design and performance of the proteomic studies.

Funding for the project was obtained from The Swedish Research Council (dnr: KT), The Swedish Brain Foundation (dnrs, KT) and the Committee for Research at Karolinska Institutet (consolidator grant, K.T.)

