## Supplemental figures and tables for "Transcriptomic-guided compound prioritization and proteomics validation for *HNRNPU* deficiency identify signalling correction"

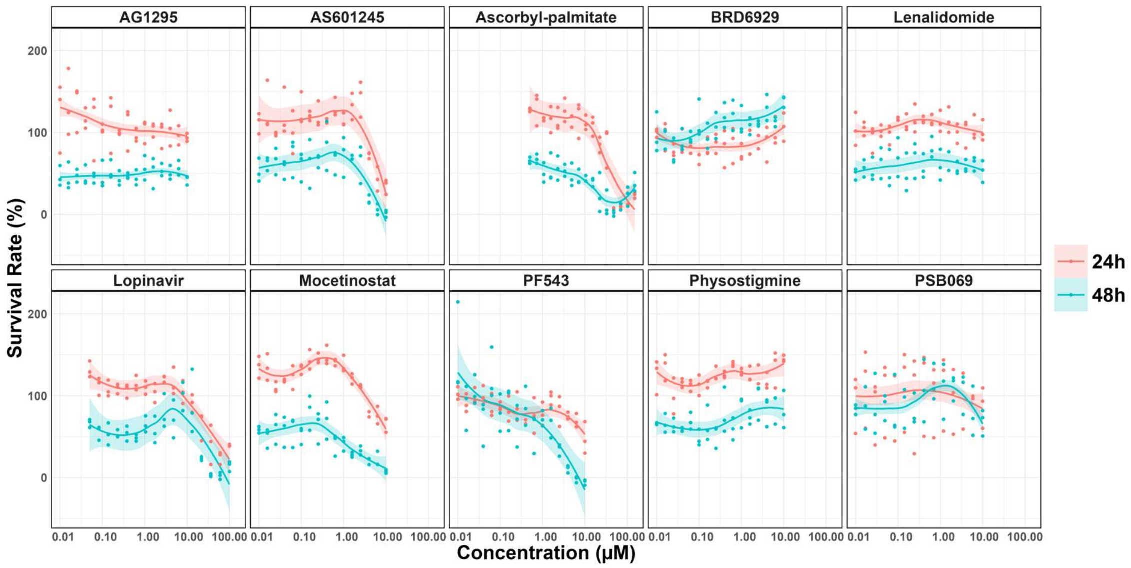

**Appendix Figure S1.** Dose-response curves illustrating the effects of prioritized compounds on cell viability in CTRL cells. Each panel represents viability data for a specific compound, with cells treated at 16 concentrations for 24 hours (pink line) or 48 hours (cyan line). Viability is presented as a percentage of vehicle (DMSO) controls. Data are presented as mean ± SEM from n = 3 technical replicates. No statistical analyses were performed.

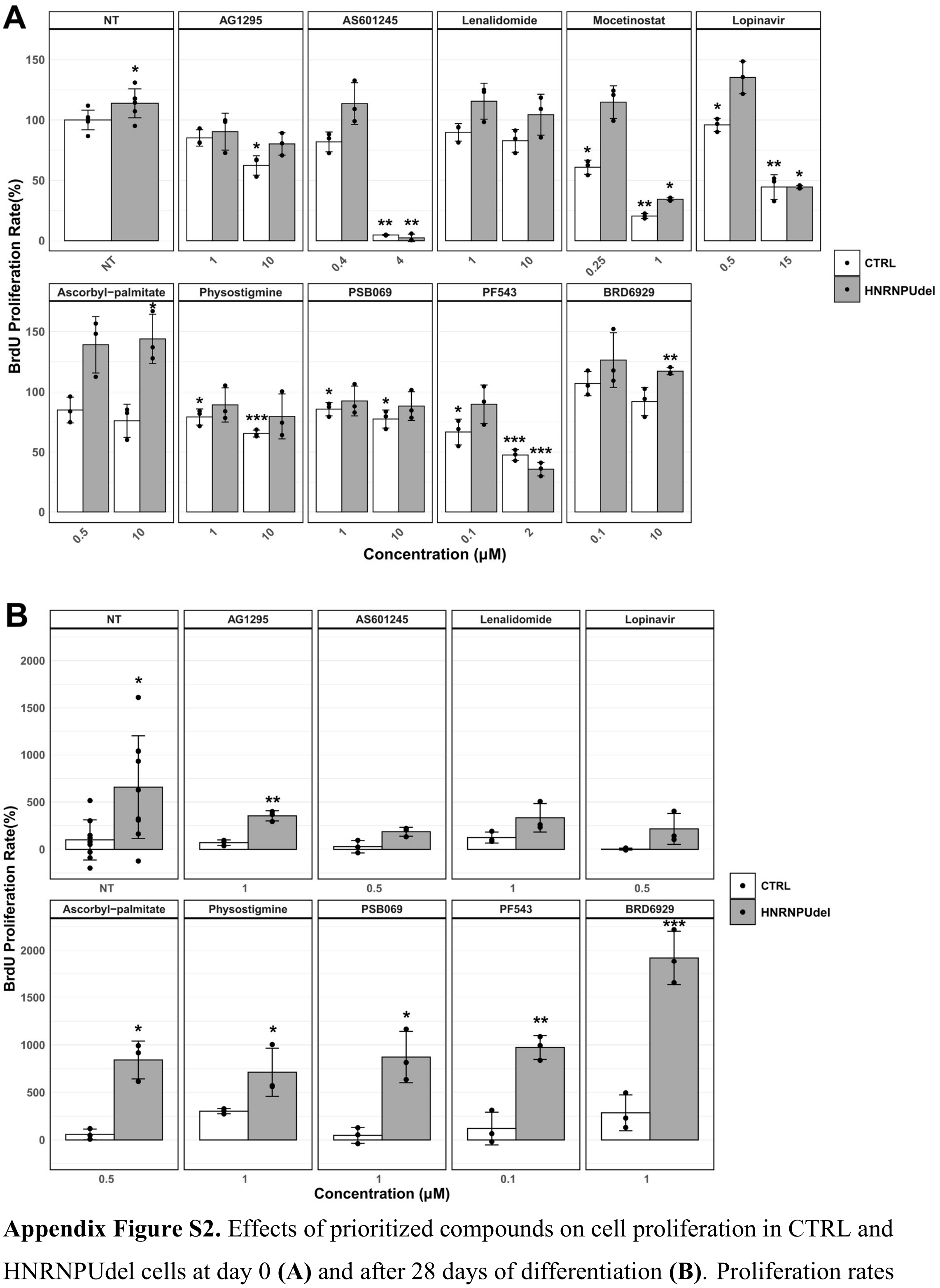

**Appendix Figure S2.** Effects of prioritized compounds on cell proliferation in CTRL and HNRNPUdel cells at day 0 **(A)** and after 28 days of differentiation **(B)**. Proliferation rates were assessed by BrdU incorporation assay and normalized to non-treated (NT) CTRL cells (100%). White bars represent CTRL cells; grey bars represent HNRNPUdel cells. Data are presented as mean ± SEM (n = 3 technical replicates). Statistical significance compared to CTRL cells: *$p < 0.05$, **$p < 0.01$, ***$p < 0.001$ (Student's t-test).

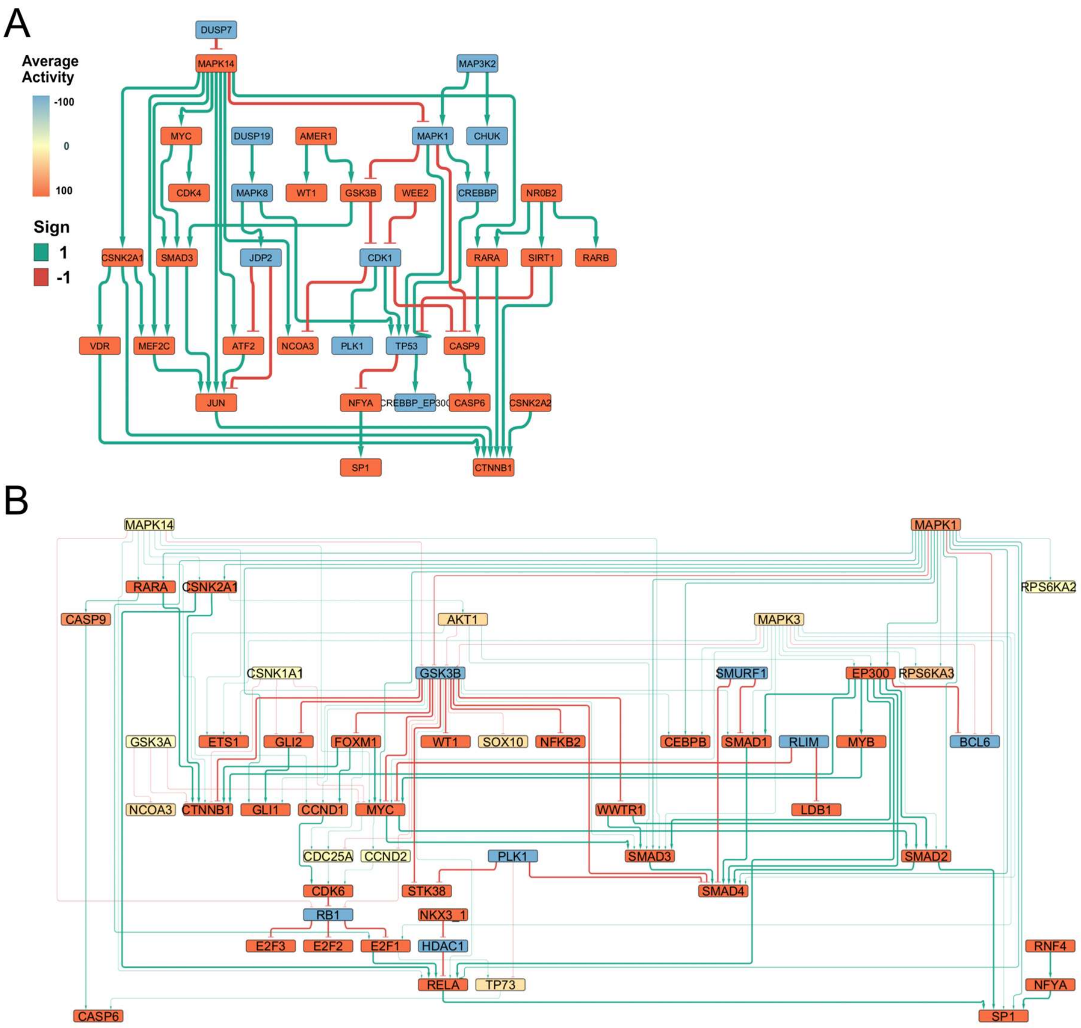

**Appendix Figure S3.** Inferred causal networks from D0 **(A)** and D28 **(B)** transcriptomic signatures in HNRNPUdel cells. All nodes are included, with a betweenness centrality > 0.

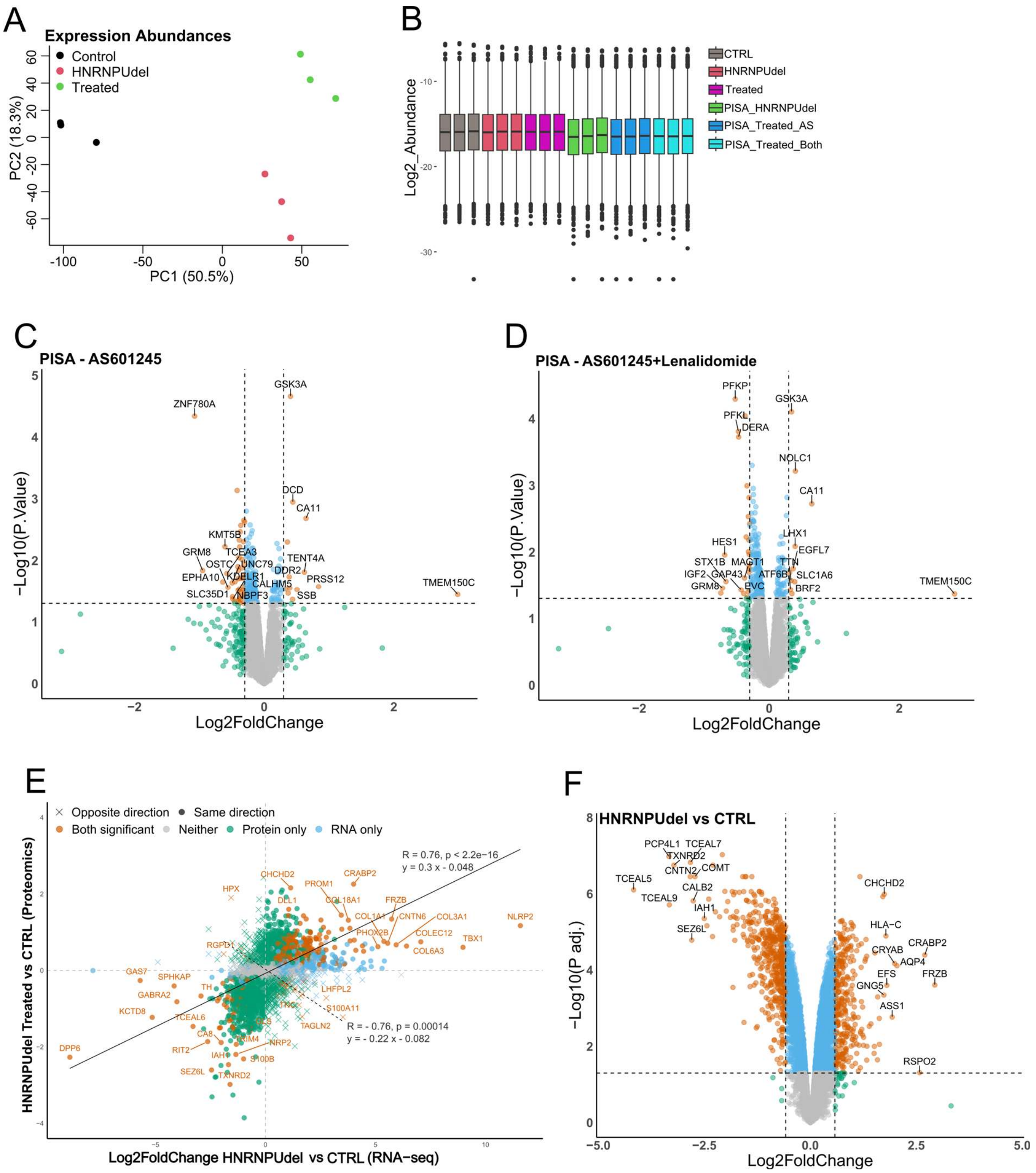

**Appendix Figure S4. (A)** PCA of normalized protein abundances of global expression changes across CTRL, HNRNPUdel non-treated (NT), and HNRNPUdel **Comb1**-treated.

**(B)** Boxplot of Log2 transformed normalized abundances for each group in global proteomics and PISA analysis.

**(C-D)** Volcano plots of protein expression changes of PISA assays after single AS601245 treatment **(C)** and combined drug treatment **(D)**. Points are coloured by significance (Significance thresholds: unadjusted p value < 0.05, |Log2(Fold Change)| > 0.25).

**(E)** Scatter plot of matched Log2(Fold Change) values comparing RNA-sequencing (CTRL versus HNRNPUdel) and proteomics data (CTRL versus HNRNPUdel combined drug treated) at D28. Points represent individual genes/proteins and colours represent the significance (orange: significant in both datasets; dark green: significant only in proteomics; sky blue: significant only in RNA-sequencing; grey: significant in neither dataset). Correlation curves of the same directional changed points (dots and solid line) and opposite directional changes (crosses and dashed line) and correlation coefficients are shown.

**(F)** Volcano plot depicting protein expression changes in HNRNPUdel versus CTRL. Points are coloured by significance (significance thresholds: adjusted p value < 0.05, |Log2(Fold Change)| > 0.58).

### Appendix Table S1. Compound comparison between transcriptomic datasets from *HNRNPU* deficient human and mouse models

| Name | Description | Direction D28 | Direction_dugger hippo | Direc_Dugger cortical | Direc_Ressler | Direc_Sapir | HNRNPUdel vs CTRL D5 |
|---|---|---|---|---|---|---|---|
| treprostinil | Prostacyclin analog | up | #N/A | #N/A | #N/A | #N/A | up |
| tyrphostin-AG-1295 | PDGFR receptor inhibitor | up | #N/A | #N/A | up | up | up |
| VU-0420363-1 | SARS coronavirus 3C-like protease inhibitor | up | #N/A | #N/A | #N/A | up | up |
| CG-930 | JNK inhibitor | up | #N/A | #N/A | #N/A | up | up |
| SU-11274 | Hepatocyte growth factor receptor inhibitor | up | #N/A | #N/A | up | #N/A | #N/A |
| tubacin | HDAC inhibitor | up | #N/A | #N/A | down | #N/A | up |
| Lenalidomide | Antineoplastic | up | up | #N/A | up | up | up |
| etamivan | Respiratory stimulant | up | #N/A | #N/A | up | up | up |
| sitagliptin | Dipeptidyl peptidase inhibitor | up | #N/A | #N/A | down | up | up |
| LM-1685 | Cyclooxygenase inhibitor | up | #N/A | #N/A | #N/A | #N/A | up |
| BAS-09104376 | HIV integrase inhibitor | up | #N/A | #N/A | up | up | up |
| iloperidone | Dopamine receptor antagonist | up | #N/A | #N/A | #N/A | #N/A | up |
| benzohydroxamic-acid | Antifungal | up | down | down | down | #N/A | #N/A |
| synephrine | Adrenergic receptor agonist | up | #N/A | #N/A | #N/A | #N/A | up |
| IB-MECA | Adenosine receptor agonist | up | #N/A | #N/A | down | #N/A | up |
| L-368899 | Oxytocin receptor antagonist | up | #N/A | #N/A | #N/A | up | #N/A |
| sinensetin | Cyclooxygenase inhibitor | up | #N/A | #N/A | #N/A | #N/A | up |
| ascorbyl-palmitate | Antioxidant | up | #N/A | #N/A | down | up | up |
| naltrexone | Opioid receptor antagonist | up | #N/A | #N/A | up | #N/A | #N/A |
| LY-303511 | Casein kinase inhibitor | up | #N/A | down | #N/A | #N/A | up |
| orantinib | FGFR inhibitor | up | #N/A | #N/A | up | up | up |
| GSK-429286A | Rho associated kinase inhibitor | up | up | #N/A | down | up | up |
| 4-(2-Amino-ethyl)-benzenesulfonamide | carbonic anhydrase inhibitor | up | #N/A | #N/A | #N/A | up | up |
| AG-370 | PDGFR receptor inhibitor | up | #N/A | #N/A | #N/A | #N/A | up |
| tocainide | Sodium channel blocker | up | #N/A | #N/A | #N/A | #N/A | up |
| griseofulvin | Tubulin inhibitor | up | #N/A | #N/A | up | up | up |
| gliquidone | Sulfonylurea | up | #N/A | #N/A | #N/A | #N/A | up |
| AT-9283 | JAK inhibitor | up | #N/A | #N/A | down | #N/A | up |
| nicergoline | Adrenergic receptor antagonist | up | #N/A | #N/A | down | #N/A | up |
| AC-55649 | Retinoid receptor agonist | up | #N/A | #N/A | #N/A | up | up |
| lacidipine | Calcium channel blocker | up | #N/A | #N/A | #N/A | #N/A | #N/A |
| leu-enkephalin | Opioid receptor agonist | up | #N/A | #N/A | #N/A | #N/A | up |
| lupanine | Sodium channel blocker | up | #N/A | #N/A | down | #N/A | up |
| danazol | Estrogen receptor antagonist | up | #N/A | #N/A | #N/A | #N/A | up |
| gamma-homolinolenic-acid | Cholesterol inhibitor | up | #N/A | #N/A | #N/A | #N/A | up |
| dovitinib | EGFR inhibitor | up | #N/A | #N/A | down | #N/A | #N/A |
| nikkomycin | Chitin inhibitor | up | #N/A | #N/A | down | up | up |
| oxymetholone | Androgen receptor agonist | up | up | #N/A | up | up | #N/A |
| clofazimine | GK0582 inhibitor | up | #N/A | down | up | up | #N/A |
| pindolol | Adrenergic receptor antagonist | up | #N/A | #N/A | down | up | up |
| linezolid | Bacterial 50S ribosomal subunit inhibitor | up | #N/A | #N/A | #N/A | up | #N/A |
| beclometasone | Glucocorticoid receptor agonist | up | #N/A | #N/A | down | up | up |
| CO-101244 | Ionotropic glutamate receptor antagonist | up | #N/A | #N/A | #N/A | #N/A | #N/A |
| carbinoxamine | Histamine receptor antagonist | up | #N/A | #N/A | down | up | up |
| desoxycortone | Mineralocorticoid receptor agonist | up | #N/A | #N/A | down | up | up |
| irinotecan | Topoisomerase inhibitor | up | #N/A | #N/A | #N/A | #N/A | #N/A |
| eicosatetraynoic-acid | Cyclooxygenase inhibitor | up | #N/A | #N/A | #N/A | #N/A | up |
| terbutaline | Adrenergic receptor agonist | up | #N/A | #N/A | #N/A | up | up |
| danusertib | Aurora kinase inhibitor | up | #N/A | #N/A | #N/A | #N/A | #N/A |
| kinetin-riboside | Apoptosis stimulant | up | #N/A | #N/A | up | up | up |
| noscapine | Bradykinin receptor antagonist | up | #N/A | #N/A | down | up | up |
| lopinavir | HIV protease inhibitor | up | #N/A | #N/A | up | up | up |
| BRD-K88742110 | HDAC inhibitor | up | up | down | up | up | #N/A |
| GSK-461364 | PLK inhibitor | up | up | #N/A | #N/A | #N/A | #N/A |
| flucytosine | Antifungal | up | #N/A | #N/A | down | up | #N/A |
| melperone | Serotonin receptor antagonist | up | #N/A | #N/A | #N/A | up | up |

| | | | | | | | |
|---|---|---|---|---|---|---|---|
| rucaparib | PARP inhibitor | up | #N/A | #N/A | down | #N/A | #N/A |
| betamethasone | Glucocorticoid receptor agonist | up | #N/A | #N/A | down | up | up |
| ALW-II-49-7 | Ephrin inhibitor | up | #N/A | #N/A | down | up | up |
| ezetimibe | Niemann-Pick C1-like 1 protein antagonist | down | #N/A | #N/A | #N/A | #N/A | down |
| salbutamol | Adrenergic receptor agonist | down | #N/A | #N/A | #N/A | #N/A | down |
| gelsemine | Acetylcholine receptor antagonist | down | #N/A | #N/A | #N/A | #N/A | down |
| sulmazole | Adenosine receptor antagonist | down | #N/A | #N/A | #N/A | #N/A | down |
| AM-404 | Cyclooxygenase inhibitor | down | #N/A | #N/A | #N/A | #N/A | down |
| RG-14620 | EGFR inhibitor | down | #N/A | #N/A | #N/A | #N/A | #N/A |
| CGS-15943 | Adenosine receptor antagonist | down | #N/A | #N/A | #N/A | #N/A | down |
| PSB-069 | NTPDase inhibitor | down | down | #N/A | #N/A | down | down |
| OBAA | Phospholipase inhibitor | down | #N/A | #N/A | #N/A | #N/A | down |
| RO-90-7501 | Beta amyloid inhibitor | down | #N/A | up | #N/A | down | down |
| physostigmine | Acetylcholinesterase inhibitor | down | #N/A | down | down | down | down |
| PF-543 | Sphingosine kinase inhibitor | down | #N/A | #N/A | down | down | down |
| modafinil | Adrenergic receptor agonist | down | up | #N/A | #N/A | #N/A | down |
| palbociclib | CDK inhibitor | down | #N/A | up | #N/A | #N/A | down |
| Merck60 | HDAC inhibitor | down | down | down | #N/A | down | down |
| HDAC3-selective | HDAC inhibitor | down | down | #N/A | #N/A | #N/A | down |
| ganciclovir | DNA polymerase inhibitor | down | #N/A | up | #N/A | #N/A | down |
| cinanserin | Serotonin receptor antagonist | down | #N/A | #N/A | #N/A | #N/A | #N/A |
| HG-5-113-01 | Protein kinase inhibitor | down | down | up | #N/A | #N/A | #N/A |
| AS-601245 | JNK inhibitor | down | down | down | #N/A | #N/A | down |
| ER-27319 | Mediator release inhibitor | down | #N/A | up | #N/A | #N/A | down |
| SU-11652 | Tyrosine kinase inhibitor | down | down | #N/A | #N/A | down | down |
| PHA-793887 | CDK inhibitor | down | #N/A | down | #N/A | #N/A | down |
| calcipotriol | Vitamin D receptor agonist | down | #N/A | #N/A | #N/A | #N/A | #N/A |
| BMS-345541 | IKK inhibitor | down | down | #N/A | #N/A | #N/A | down |
| JNJ-7706621 | CDK inhibitor | down | #N/A | #N/A | #N/A | #N/A | down |
| PF-562271 | Focal adhesion kinase inhibitor | down | #N/A | down | #N/A | #N/A | down |
| lestaurtinib | FLT3 inhibitor | down | #N/A | #N/A | #N/A | #N/A | down |
| moclobemide | Monoamine oxidase inhibitor | down | #N/A | #N/A | #N/A | #N/A | down |
| canrenoic-acid | Mineralocorticoid receptor antagonist | down | #N/A | up | #N/A | #N/A | down |
| camptothecin | Topoisomerase inhibitor | down | #N/A | #N/A | #N/A | #N/A | down |
| SB-205607 | Delta 1 opioid receptor agonist | down | #N/A | #N/A | #N/A | #N/A | down |
| mocetinostat | HDAC inhibitor | down | down | #N/A | #N/A | down | down |
| bisindolylmaleimide-ix | CDK inhibitor | down | #N/A | up | #N/A | #N/A | down |
| ceramide | Phosphoenolpyruvate carboxylase activator | down | #N/A | #N/A | #N/A | #N/A | down |
| 5-iodotubercidin | Adenosine kinase inhibitor | down | #N/A | up | #N/A | #N/A | down |
| flavanone | 11-beta-HSD1 inhibitor | down | #N/A | #N/A | #N/A | down | down |
| doxorubicin | Topoisomerase inhibitor | down | #N/A | #N/A | #N/A | #N/A | down |
| topotecan | Topoisomerase inhibitor | down | #N/A | up | #N/A | down | down |
| ochratoxin-a | Phenylalanyl tRNA synthetase inhibitor | down | down | down | #N/A | #N/A | down |
| pidorubicine | Topoisomerase inhibitor | down | #N/A | #N/A | #N/A | #N/A | down |
| mitoxantrone | Topoisomerase inhibitor | down | #N/A | up | #N/A | #N/A | down |
| triptolide | RNA polymerase inhibitor | down | #N/A | #N/A | #N/A | #N/A | down |
| ZG-10 | JNK inhibitor | down | #N/A | down | #N/A | #N/A | down |
| SD-169 | p38 MAPK inhibitor | down | #N/A | #N/A | #N/A | #N/A | #N/A |
| AG-14361 | PARP inhibitor | down | #N/A | #N/A | down | #N/A | down |
| alvocidib | CDK inhibitor | down | #N/A | #N/A | #N/A | #N/A | down |
| honokiol | AKT inhibitor | down | #N/A | #N/A | #N/A | #N/A | down |
| JAK3-inhibitor-VI | JAK inhibitor | down | down | #N/A | #N/A | #N/A | down |
| cotinine | Nicotine metabolite | down | #N/A | #N/A | #N/A | #N/A | #N/A |
| saracatinib | SRC inhibitor | down | #N/A | up | #N/A | down | down |
| pirarubicin | Topoisomerase inhibitor | down | #N/A | #N/A | #N/A | #N/A | down |
| A-443644 | AKT inhibitor | down | #N/A | up | #N/A | #N/A | down |
| ticlopidine | Purinergic receptor antagonist | down | #N/A | #N/A | #N/A | #N/A | #N/A |
| CAY-10470 | NFkB pathway inhibitor | down | #N/A | #N/A | #N/A | down | #N/A |
| PIK-75 | DNA protein kinase inhibitor | down | #N/A | #N/A | #N/A | #N/A | down |
| dorsomorphin | AMPK inhibitor | down | down | #N/A | #N/A | #N/A | down |
| gitoxigenin | ATPase inhibitor | down | #N/A | up | #N/A | #N/A | down |

| | | | | | | | |
|---|---|---|---|---|---|---|---|
| cyclopamine | Smoothened receptor antagonist | down | #N/A | #N/A | #N/A | #N/A | down |
| PI-103 | MTOR inhibitor | down | #N/A | up | #N/A | #N/A | down |
| epinephrine | carbonic anhydrase activator | down | #N/A | #N/A | #N/A | #N/A | #N/A |
| targinine | Nitric oxide synthase inhibitor | down | #N/A | #N/A | #N/A | #N/A | down |
| n-formylmethionylalanine | macrophage activator | down | #N/A | #N/A | #N/A | #N/A | down |
| SR-57227A | Serotonin receptor agonist | down | down | #N/A | #N/A | down | #N/A |
| rhamnetin | HDAC inhibitor | down | #N/A | #N/A | #N/A | #N/A | down |
| indapamide | Thiazide diuretic | down | #N/A | #N/A | #N/A | #N/A | #N/A |
| T-0156 | Phosphodiesterase inhibitor | down | #N/A | up | #N/A | down | down |
| bumetanide | Solute carrier family member inhibitor | down | #N/A | #N/A | #N/A | #N/A | down |
| AT-7519 | CDK inhibitor | down | #N/A | #N/A | #N/A | #N/A | down |
| berbamine | Calmodulin antagonist | down | down | #N/A | #N/A | down | #N/A |
| linifanib | PDGFR receptor inhibitor | down | down | #N/A | #N/A | down | down |
| SB-218078 | CHK inhibitor | down | #N/A | #N/A | up | #N/A | down |
| dehydrocholic-acid | choleretic agent | down | #N/A | #N/A | #N/A | #N/A | #N/A |
| ascorbic-acid | Antioxidant | down | #N/A | #N/A | #N/A | down | down |
| nicorandil | Nitric oxide donor | down | #N/A | #N/A | #N/A | #N/A | #N/A |
| piretanide | Glucocorticoid receptor agonist | down | #N/A | up | #N/A | #N/A | down |
| CGP-57380 | MAP kinase inhibitor | down | #N/A | up | #N/A | #N/A | down |
| entinostat | HDAC inhibitor | down | down | down | #N/A | down | down |
| bisbenzimide | DNA binding agent | down | #N/A | #N/A | #N/A | #N/A | down |
| CAY-10578 | Casein kinase inhibitor | down | #N/A | #N/A | #N/A | down | #N/A |
| hinokitiol | Tyrosinase inhibitor | down | #N/A | #N/A | #N/A | down | #N/A |
| erythromycin | NFkB pathway inhibitor | down | #N/A | #N/A | #N/A | down | down |
| methandriol | Androgenic steroid | down | #N/A | #N/A | #N/A | #N/A | #N/A |
| staurosporine | PKC inhibitor | down | #N/A | #N/A | #N/A | #N/A | down |
| piperlongumine | Glutathione transferase inhibitor | down | down | #N/A | #N/A | down | down |
| nonoxynol-9 | Membrane integrity inhibitor | down | #N/A | up | #N/A | #N/A | down |
| LY-16350 | Dopamine receptor agonist | down | #N/A | up | #N/A | #N/A | down |
| dihydroergocristine | Adrenergic receptor antagonist | down | down | up | #N/A | down | down |
| gamma-linolenic-acid | Cyclooxygenase inhibitor | down | #N/A | up | #N/A | #N/A | down |
| idarubicin | Topoisomerase inhibitor | down | #N/A | #N/A | #N/A | #N/A | down |
| zoxazolamine | Myorelaxant | down | #N/A | #N/A | #N/A | #N/A | down |
| parachlorophenol | Anti-infective | down | #N/A | up | #N/A | down | down |
| famotidine | Histamine receptor antagonist | down | #N/A | #N/A | #N/A | #N/A | down |
| FK-888 | Tachykinin antagonist | down | #N/A | #N/A | #N/A | #N/A | #N/A |
| tacedinaline | HDAC inhibitor | down | #N/A | #N/A | #N/A | #N/A | down |
| ryuvidine | Histone lysine methyltransferase inhibitor | down | down | #N/A | #N/A | down | down |
| diflunisal | Prostanoid receptor antagonist | down | #N/A | #N/A | #N/A | #N/A | down |
| SN-38 | Topoisomerase inhibitor | down | #N/A | #N/A | #N/A | #N/A | down |
| LY-83583 | Guanylyl cyclase inhibitor | down | #N/A | #N/A | #N/A | #N/A | down |
| asiatic-acid | Apoptosis stimulant | down | #N/A | #N/A | #N/A | #N/A | #N/A |
| strophanthidin | ATPase inhibitor | down | #N/A | up | #N/A | down | #N/A |
| EMF-bca1-60 | caspase inhibitor | down | up | #N/A | #N/A | down | down |
| dactinomycin | RNA polymerase inhibitor | down | #N/A | up | #N/A | #N/A | down |
| NCH-51 | HDAC inhibitor | down | #N/A | #N/A | #N/A | down | down |
| milnacipran | Serotonin reuptake inhibitor | down | #N/A | #N/A | #N/A | #N/A | #N/A |
